# Neuronal activity and amyloid-β cause tau seeding in the entorhinal cortex in Alzheimer’s disease

**DOI:** 10.1101/2025.03.07.642054

**Authors:** Christoffer G. Alexandersen, Dani S. Bassett, Alain Goriely, Pavanjit Chaggar, the Alzheimer’s Disease Neuroimaging Initiative

## Abstract

The entorhinal cortex is the earliest site of tau pathology in both Alzheimer’s disease and primary age-related tauopathy, yet the mechanisms underlying this selective vulnerability remain poorly understood. Here, we use a computational model integrating neuronal activity and amyloid-*β* deposition with interneuronal tau transport to predict regional susceptibility to tau seeding. Using fluorodeoxyglucose PET as a measure of neuronal activity, we show that brain-wide activity patterns drive tau accumulation in the medial temporal lobe, independent of amyloid status. Incorporating amyloid PET, we further show that amyloid-*β* selectively amplifies tau seeding in the entorhinal cortex, aligning with its early involvement in Alzheimer’s disease. These predictions are supported by cross-subject correlation analysis, which reveals a significant association between model-derived seeding concentrations and empirical tau deposition. Our findings suggest that neuronal activity patterns shape the early landscape of tau pathology, while amyloid-*β* deposition creates a unique vulnerability in the entorhinal cortex, potentially triggering the pathological cascade that defines Alzheimer’s disease.

## 1. Introduction

Tau pathology is a defining characteristic of Alzheimer’s disease (AD) and primary age-related tauopathy (PART), with the medial temporal lobe (MTL) typically being the first region to exhibit tau accumulation, particularly in the entorhinal cortex (EC).^1–5^ Despite consistent evidence demonstrating tau pathology in the EC during the initial stages of AD, we currently have a poor understanding of *why* the EC is the initial site of tau pathology in AD. Gaining such an understanding may be crucial for the development of early interventions to prevent or delay tau-related pathology in AD.

Toxic tau is believed to propagate across the brain primarily via cell-to-cell transmission; however, the mechanisms through which this transport process is regulated are unclear. Experiments in transgenic animal models have demonstrated that misfolded tau proteins, either injected into the brain or over-expressed in the EC, result in local tau accumulation and propagation to nearby regions.^6–10^ This evidence is complemented by studies using intracranial injections of synthetic tau fibrils or human brain-derived pathological tau into mouse models to show that tau aggregates can propagate through axonal connections.^11–14^ This process has been further validated using human neuroimaging studies, in which models describing trans-synaptic spread of tau pathology accurately predict tau progression.^15–17^ However, other factors such as neuronal activity and amyloid-*β* (A*β*) also modulate the tau transport process. *In vitro* studies have shown that greater neuronal activity results in increased interneuronal tau transport,^18–20^ and animal models combining A*β* and tau pathology show enhanced propagation when A*β* pathology is present.^21,22^ These mechanisms may indeed be linked, since A*β* has been shown to induce neuronal hyperexcitability in rat slices and *in vivo* mouse models,^23–25^ suggesting that A*β* may exert its influence on tau spreading through neuronal hyperexcitability. Further confounding the relationship between A*β* and tau is their spatial properties—while tau pathology in PART and AD begins in the EC,^1–3,5,26^ early A*β* depositions are mainly present in the neocortex,^26,27^ suggesting that the effects of A*β* on tau may be non-local. Overall, the relationship between A*β*, tau, and neuronal activity is complex,^28^ and it remains unclear whether A*β* deposition directly enhances tau seeding or if it modulates tau spread through other means, such as A*β*-induced hyperexcitability. Particularly puzzling is the fact that tau accumulation occurs in the MTL without the involvement of A*β* deposition in PART.^1–3,5^ Thus, A*β* deposition is *not* necessary for early tau accumulation in the MTL, but *is* necessary for tau to spread from the MTL to neocortical regions

In a recent analysis of a mathematical model coupling neuronal activity with interneuronal protein transport, we showed that neuronal activity gradients promote the localized seeding and aggregation of tau protein in regions of low relative activity.^29^ This finding suggests that brain-wide activity gradients may explain the regional susceptibility to tau seeding. Here, we use a simplified version of the model introduced in Ref.^29^ and integrate it with empirical data to predict which brain regions are most susceptible to early tau seeding, based on the hypothesis that tau axonal transport is accelerated by either (i) neuronal activity or (ii) A*β* deposition. Using FDG PET imaging as a measure of neuronal activity, we demonstrate that brain-wide activity patterns drive tau seeding in the medial temporal lobe. The seeding of tau in the medial temporal lobe is consistent across subject classifications based on A*β* and tau levels. Furthermore, under the assumption that A*β* increases tau transfer, we show that A*β* deposition patterns predict the entorhinal cortex to be the most vulnerable region to tau seeding, with a particularly strong bias towards entorhinal seeding for early-stage Alzheimer’s subjects. We also show that the model predictions of early Braak stage regions as tau seeding regions are statistically significant upon re-shuffling PET signals by region for the FDG- and A*β*-based seeding predictions. Finally, subject-level analysis confirms that predicted seeding concentrations correlate significantly with empirical tau deposition, supporting the hypothesis that neuronal activity and amyloid deposition shape early tau pathology.

The modeling predictions and validation support the hypothesis that brain-wide gradients of neuronal activity lead to a greater accumulation of tau seeds in the medial temporal lobe, consistent with the observation that tau pathology accumulates in this region even in elderly individuals without A*β* deposition, as seen in PART. Additionally, our results suggest that A*β* deposition leads to a particularly high accumulation of tau in the entorhinal cortex, which is consistently the first region to accumulate tau in Alzheimer’s disease. Together, these findings suggest that brain-wide neuronal activity patterns determine the susceptibility of the medial temporal lobe to tau pathology and that A*β* deposition amplifies tau accumulation specifically in the entorhinal cortex, providing a compelling explanation for why Alzheimer’s disease initially manifests in this region and potentially triggers the pathological cascade that spreads throughout the brain.

## 2. Materials & Methods

### 2.1. Data processing

We use PET from the Alzheimer’s Disease Neuroimaging Initiative (ADNI, adni.loni.usc.edu). ADNI is a public-private partnership with the aim of using serial biomarkers to measure the progression of AD. For up-to-date information, see www.adni-info.org. For A*β* and tau, we download the fully processed PET tabular data, summarized as SUVR values on the Desikan-Killiany (DK) atlas.^30^ For each subject, the first pair of A*β* and tau PET scans was identified to determine the subject’s A*β* and tau status. A*β* SUVR was calculated using the composite reference region provided by ADNI; then, A*β* status was classified as A*β*-positive (A*β*^+^) if the ADNI summary composite region SUVR *>* 0.78 and A*β*-negative (A*β*^*−*^) otherwise. Tau-PET SUVR was calculated using the inferior cerebellar SUVR as a reference region. Tau status was stratified into three categories based on the SUVR in two composite regions: the MTL, comprising the bilateral entorhinal cortices and amygdalae, and a neocortical area, comprising the bilateral inferior and middle lateral temporal lobes. The three tau categories are then: MTL tau-negative (*τ* MTL^*−*^), MTL tau-positive (*τ* MTL^+^), and MTL and neocortical tau-positive (*τ* NEO^+^) categories. The threshold for tau composite SUVR using an inferior cerebellum reference are 1.375 and 1.395 for the MTL and cortical composites, respectively, and are derived using regional Gaussian mixture modeling (see^31^ for details). For A*β* PET scans, we extracted in total 1642 scans (870 scans in A*β*^*−*^, 185 scans in A*β*^+^*τ* MTL^*−*^*τ* NEO^*−*^, 61 scans in A*β*^+^*τ* MTL^+^*τ* NEO^*−*^, and 152 scans in A*β*^+^*τ* MTL^+^*τ* NEO^+^). Demographic characteristics of the A*β* PET study participants are summarized in Table 1.

**Table 1:**
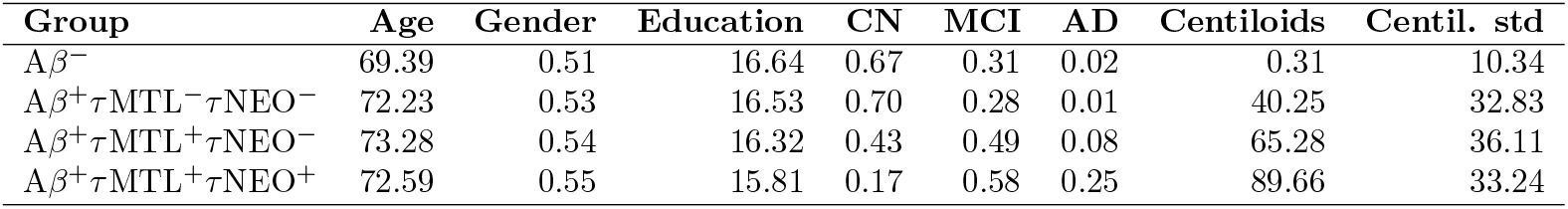
Demographic and clinical characteristics of A*β* PET study participants. Mean age, gender distribution (proportion of females), years of education, proportions of cognitively normal (CN), mild cognitive impairment (MCI), Alzheimer’s disease (AD), and centiloid means and standard deviations across biomarker-defined groups

| Group | Age | Gender | Education | CN | MCI | AD | Centiloids | Centil. std |
| --- | --- | --- | --- | --- | --- | --- | --- | --- |
| A$\beta^-$ | 69.39 | 0.51 | 16.64 | 0.67 | 0.31 | 0.02 | 0.31 | 10.34 |
| A$\beta^+$ τMTL$^-$ τNEO$^-$ | 72.23 | 0.53 | 16.53 | 0.70 | 0.28 | 0.01 | 40.25 | 32.83 |
| A$\beta^+$ τMTL$^+$ τNEO$^-$ | 73.28 | 0.54 | 16.32 | 0.43 | 0.49 | 0.08 | 65.28 | 36.11 |
| A$\beta^+$ τMTL$^+$ τNEO$^+$ | 72.59 | 0.55 | 15.81 | 0.17 | 0.58 | 0.25 | 89.66 | 33.24 |

To classify FDG scans based on their A*β* and tau status, we collect a subject’s first FDG scans taken within one year of an A*β* and tau PET scan. Subjects without an FDG scan meeting this temporal criterion were excluded from further analysis. FDG PET SUVR was calculated using the pons as a reference region, which is minimally affected by AD.^32,33^ For the FDG PET scans, we extracted in total 321 scans (138 scans in A*β*^*−*^, 48 scans in A*β*^+^*τ* MTL^*−*^*τ* NEO^*−*^, 39 scans in A*β*^+^*τ* MTL^+^*τ* NEO^*−*^, and 90 scans in A*β*^+^*τ* MTL^+^*τ* NEO^+^). Demographic characteristics of the FDG PET study participants are summarized in Table 2.

**Table 2:** Demographic and clinical characteristics of FDG PET study participants. Mean age, gender distribution (proportion of females), years of education, and proportions of cognitively normal (CN), mild cognitive impairment (MCI), Alzheimer’s disease (AD), and centiloid means and standard deviations across biomarker-defined groups.

| Group | Age | Gender | Education | CN | MCI | AD | Centiloids | Cent. std |
| --- | --- | --- | --- | --- | --- | --- | --- | --- |
| A$\beta^-$ | 70.58 | 0.37 | 16.23 | 0.09 | 0.80 | 0.10 | -1.58 | 11.71 |
| A$\beta^+$ τMTL$^-$ τNEO$^-$ | 73.84 | 0.42 | 15.96 | 0.17 | 0.79 | 0.04 | 51.50 | 34.49 |
| A$\beta^+$ τMTL$^+$ τNEO$^-$ | 74.21 | 0.44 | 16.79 | 0.05 | 0.79 | 0.15 | 72.38 | 37.31 |
| A$\beta^+$ τMTL$^+$ τNEO$^+$ | 71.77 | 0.52 | 15.60 | 0.01 | 0.63 | 0.36 | 98.22 | 31.97 |

We downloaded the maximally preprocessed image data provided by ADNI. Images were coregistered to the MNI152 standard space Advanced Normalization Tools (ANTs).^34^ We use the DK atlas for regional parcellation and signal summarization to be consistent with preprocessed A*β* PET data from ADNI. Once normalized to the reference region, the mean SUVR is calculated for each DK region. For both A*β* PET and FDG PET, we correct for spurious signal in the frontal pole, temporal pole, and banks of the superior temporal sulcus by setting the SUVR of these regions to be the average of itself and adjacent regions.^35^

### 2.2. Computational modeling

Mathematical models have been used to study the prion-like spread of proteins in various neurodegenerative diseases.^36–40^ These models simulate the spread and increase of protein pathology through structural connectomes—brain networks that map the physical architecture of the brain, inferred from diffusion tensor or weighted imaging and tractography. These computational models show promising agreement with empirical data.^17,31^ The simplest class of spreading models are linear diffusion models, which describe a transport process across a network.^36^ These linear diffusion models may be complemented with nonlinear dynamics, often representing local prion-like replication within each brain region.^38^

In this study, however, we are not modeling the progression of tau pathology throughout the entire disease but rather the diffusion of early tau seeds before disease initiation. Our goal is to predict which regions are most susceptible to early tau seeding. Since at this time, protein densities are very low, nonlinear effects do not yet play a role and linear effects dominate. Hence, we use a linear diffusion model describing the mass-conserving transport of protein seeds between regions and local production and decay of tau seeds. Moreover, we weight the axonal transport of the protein by a specific transport process, which, in our study, will either be (1) neuronal activity or (2) amyloid deposition. Since, this effect is a modification of a passive diffusion process, we refer to it as an *anomalous transport process* (generated by neuronal activity and amyloid deposition). As we are primarily interested in the impact of this anomalous transport process, we set the protein production and degradation rates to be the same across regions.

The connectome is represented by the *N* × *N* weighted adjacency matrix **W**, where each node corresponds to a brain region, and edges are weighted as *W*_*ij*_ = *n*_*ij*_, the number of fibers from region *j* to region *i*. The regular diffusive transport between different nodes (brain regions) is usually described using the graph Laplacian **L** with entries

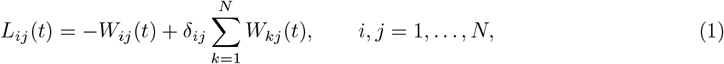

where *δ*_*ij*_ is the Kronecker symbol. In our case, however, we scale the axonal transport by an anomalous transport process. Each node has an *anomaly level* associated with it *A*_*i*_, and we collect these in a diagonal *N* × *N* matrix **A** = diag(*A*_1_, …, *A*_*n*_). We then define an anomaly-scaled adjacency matrix 𝒲 = **W**(**I** + *ε***A**), where **I** is the *N* × *N* identity matrix and *ε* the strength of the anomaly. The Laplacian ℒ built from 𝒲 is then given by

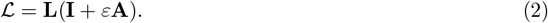

Here, we will consider a protein species **u** ∈ ℝ^*N*^, namely tau seeds, that is being naturally produced and degraded in each node (brain region) but which is also transported across edges (axonal connections), where the transport is accelerated by the anomalous transport process. This dynamical process is modeled by

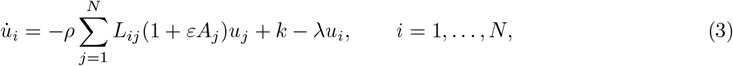

where *ρ >* 0 is the transport coefficient, *k >* 0 is the protein production rate, and *λ >* 0 is the protein degradation rate. We will assume at all times that the production and degradation rates are the same across all regions as we focus on the transport process.

Note that as *ε* → 0, we recover the traditional transport process that depends only on the structural connectivity. As *ε* increases, so does the anomalous transport process’s impact on the protein spreading. We gain more insight into the impact of the anomalous transport process by considering the eigenvector corresponding to the zero eigenvalue of the anomaly-weighted Laplacian ℒ. A graph Laplacian with a single, fully connected component always has a zero eigenvalue with a 1-dimensional eigenspace. The 0-eigenvector is, in these cases, the stationary solution (up to scaling) of 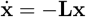. If **v** is the 0-eigenvector of the Laplacian built from the structural connectivity **L** alone, then the 0-eigenvector of the anomaly-weighted Laplacian ℒ has components *v*_*i*_*/*(1 + *εA*_*i*_). In other words, the anomalous transport process **A** leads to higher accumulation in regions with low *A*_*i*_ and lower accumulation in regions with high *A*_*i*_. Hence a region with low activity surrounded by neighbors with high activity will naturally accumulate proteins.

Here, we are interested in the impact of neuronal activity and amyloid deposition on tau seeding. Therefore, we focus on parameter regimes in which the production of tau seeds is small compared to protein transport. In such a parameter regime, the differences in tau seeding across regions will be mainly impacted by the anomalous transport process, which will either be neuronal activity or amyloid deposition. As such, the following parameters were used throughout the study: *ρ* = 0.7, *k* = 0.014, and *λ* = 0.14, whereas *ε* is used as a control parameter. All simulations and analyses were written in the Julia programming language, and the differential equations were solved using the Tsitouras 5th-order method from the DifferentialEquations Julia package with an absolute and relative tolerance of 10^*−*10^.

### 2.3. Statistical analysis

#### 2.3.1. Assessing group differences in predicted tau seeding

To determine whether tau seeding predictions differ between subjects based on A*β* and tau status, we compare the average predicted seeding concentrations between the A*β*^*−*^ group and the amyloid- or tau-positive groups. We use each individual’s FDG and A*β* PET SUVRs to predict their personalized seeding concentration in Braak stage 1 regions (bilateral EC), generating a distribution for each group. We then use two-sided Welch’s *t* − tests to assess whether the mean predicted seeding concentrations differ significantly between the amyloid-negative and the amyloid-positive/tau-positive groups. The significance levels are defined as *p <* 0.05 (*), *p <* 0.01 (**), and *p <* 0.001 (***). A Bonferroni correction was applied to adjust the significance levels for multiple comparisons to the same A*β*^*−*^ distributions. The same procedure is applied to Braak stage 2 regions.

#### 2.3.2. Assessing the significance of predicted tau seeding regions

Using our computational model, we predict seeding regions and test the significance of our predictions under randomized regional PET SUVRs. Our null hypothesis is that the particular regional levels of neuronal activity and amyloid deposition (regardless of subject classification based on amyloid and tau levels) do not contribute to tau seeding in Braak stage 1 (or stage 2) regions. The alternative hypothesis is that the neuronal activity or amyloid deposition (for some subject classification) contributes to tau seeding in Braak stage 1 (or stage 2). To evaluate the probability of observing our seeding predictions given the null hypothesis, we construct a null model with PET SUVRs shuffled across regions. Depending on the simulation, the A*β* PET or FDG PET values are shuffled randomly across all regions of interest to evaluate the probability of observing our findings. This shuffling process was repeated 10,000 times, and the concentrations of tau seeds were predicted and stored for each iteration. Predicted concentrations from the null model across all regions and trials were aggregated into a single dataset, forming a distribution of null concentrations. We then define a *seeding threshold* as the midpoint in the highest gap between null concentrations observed across all regions and trials (see Fig. S1 and S2). A region in the unshuffled simulation is deemed “seeded” if its predicted concentration exceeds this null threshold.

For statistical validation, we performed two tests. First, we counted the number of seeding regions belonging to Braak stage 1 in our non-shuffled modeling predictions. Using the null model, we calculated the probability of observing these many or more seeded regions in Braak stage 1 with randomly shuffled regional PET values. This probability served as the *p*-value for Braak stage 1 predictions. Second, the same procedure was repeated but seeded regions belonging to both Braak stages 1 and 2 were counted. The corresponding *p*-value was determined based on the likelihood of observing these many or more seeded regions in these combined Braak stages under the null model. The significance levels are defined as *p <* 0.05 (*), *p <* 0.01 (**), and *p <* 0.001 (***).

These *p*-values allow us to assess the probability that the regional patterns of neuronal activity and amyloid deposition observed in the study subjects contribute to tau seeding in Braak stages 1 and 2 under the null hypothesis that their particular regional values are unrelated to tau seeding.

#### 2.3.3. Assessing subject-level correlations between model-predicted seeding and early tau deposition

To examine the relationship between neuronal activity, amyloid deposition, and early tau seeding across subjects, we performed simple, least-squares linear regression. The explanatory variables were the asymptotic model-derived seeding concentrations based on subject-level FDG and amyloid PET patterns, and the predicted variables were subject-level empirical tau PET SUVRs, where both variables were summed over early Braak stages for each subject. The analysis included 188 subjects with both tau and FDG PET data and 409 subjects with both tau and amyloid PET data. We tested the null hypothesis that the regression slope was equal to zero (two-sided *t*-test), with significance levels defined as *p <* 0.05 (*), *p <* 0.01 (**), and *p <* 0.001 (***). Pearson correlation coefficients (r-values) were computed to quantify the strength of the association between the predicted and observed tau seeding across subjects.

## 3. Results

To investigate how neuronal activity and A*β* deposition shape regional susceptibility to tau seeding, we integrate empirical FDG PET and A*β* PET imaging data with a mathematical framework describing interneuronal tau transport. A schematic overview of our approach is shown in Fig. 1, which outlines the key steps in our modeling framework and analysis. In the following sections, we present our key findings: (1) brain-wide neuronal activity gradients predict tau seeding in the MTL regardless of A*β* status, (2) A*β* deposition selectively amplifies tau seeding in the entorhinal cortex, (3) seeding predictions in early Braak stage are statistically significant upon shuffling PET SUVRs, and (4) subject-level seeding predictions correlate with early empirical tau deposition.

**Figure 1.**
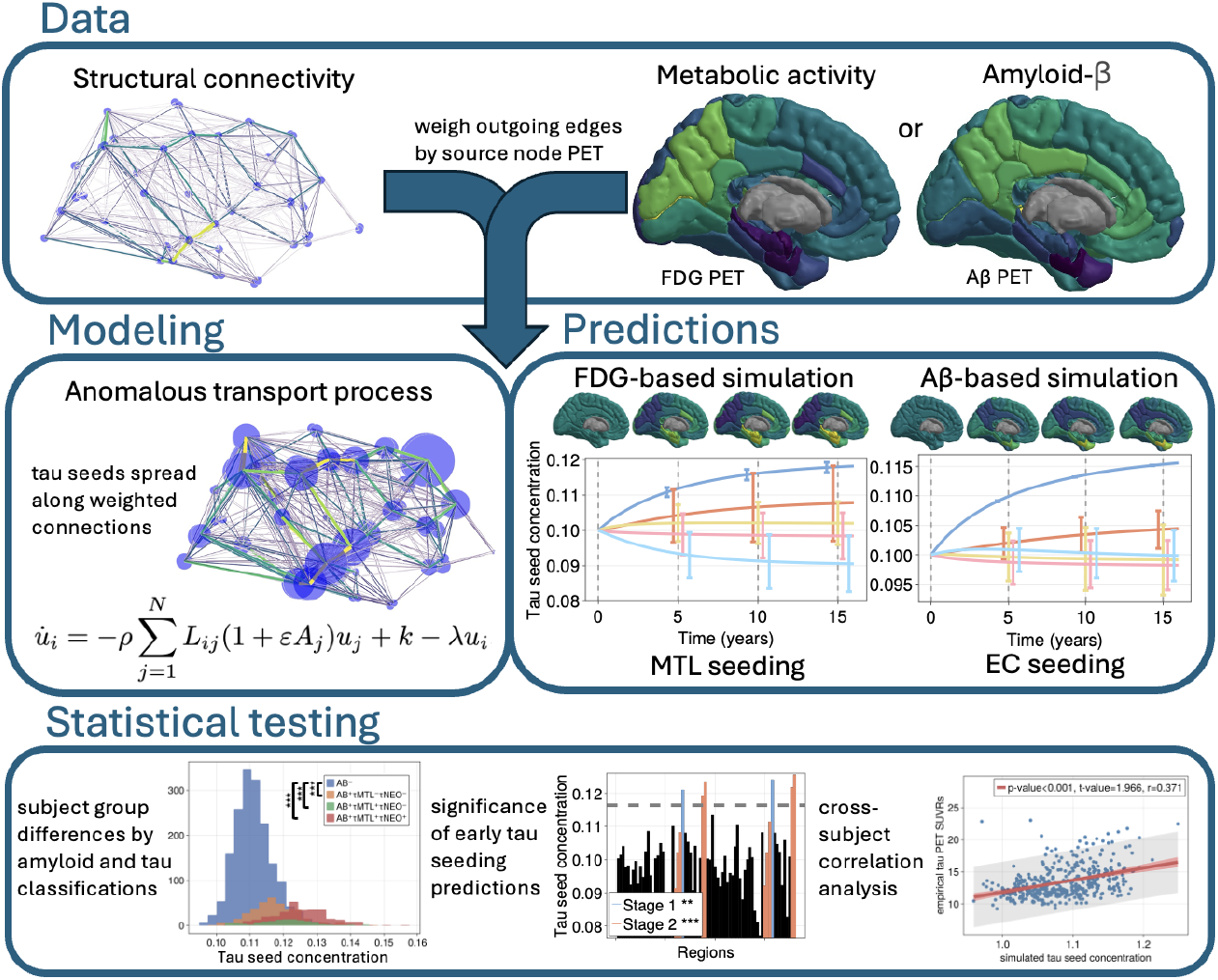
Schematic overview of our framework for modeling tau seeding susceptibility. We integrate structural connectivity with metabolic/neuronal activity (FDG PET) or amyloid-*β* deposition (A*β* PET), which are hypothesized to accelerate tau seed transport. Scaling the protein transport along structural connections by their regional PET SUVRs, we construct a mathematical model to simulate and predict regional susceptibility to tau seeding. To evaluate our predictions, we conduct three statistical tests: (1) assessing whether predicted seeding concentrations in early Braak stages differ between subject groups stratified by amyloid and tau status, (2) testing the significance of our seeding predictions by shuffling regional PET values, generating a null distribution where the spatial pattern of neuronal activity or amyloid deposition is randomized, and (3) testing whether model-derived tau seeding correlates with empirical tau accumulation across subjects.

### 3.1. Metabolic activity drives the seeding of tau in the medial temporal lobe

Many elderly people exhibit tauopathy in the medial temporal lobe, which can occur in the absence of A*β* plaques, as in PART.^5^ Recent research suggests that the transport of tau is influenced by neuronal activity.^18–20^ This evidence, in turn, suggests that neuronal activity may play a role in initiating tau pathology. Using a computational model, we simulate the spreading of tau seeds across an averaged structural connectome and use an averaged FDG PET signal to model the impact of metabolic activity on the spread. The tau seeds will spread along axonal connections but will spread faster away from regions of higher activity and towards regions of lower activity. In the model, all brain regions (nodes in the connectome) are initialized with the same number of tau seeds (**u**(*t* = 0) = 0.1), and each region produces tau seeds at the same low rate. Hence, the model has no bias towards tau accumulation in any particular region, apart from what is induced by incorporating the FDG PET signal.

As shown in Fig. 2A, using the averaged FDG PET across all subjects independent of A*β* and tau status, we observe that metabolic activity steers the tau seeds towards the medial temporal lobe in the model simulations. Specifically, the entorhinal cortex, the hippocampus, and the amygdala have a higher concentration of tau seeds than other regions.

**Figure 2.**
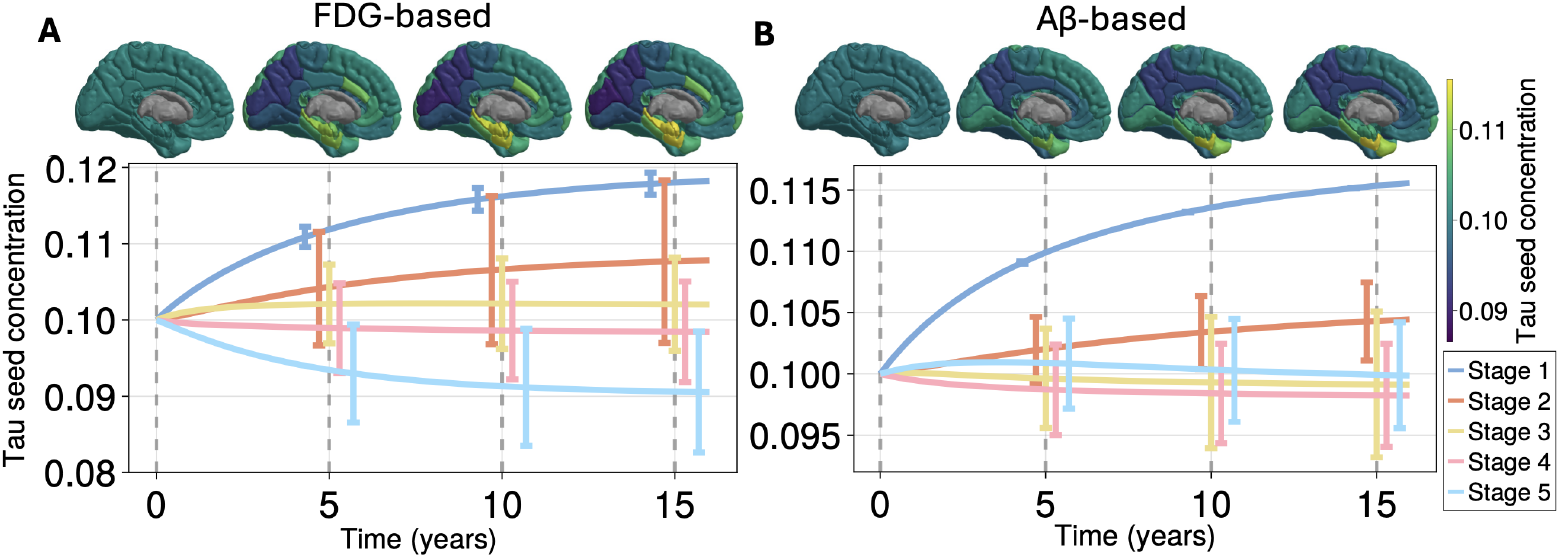
Time evolution of tau seeding concentrations across the brain in a computational model. The evolution of tau seeds in an FDG-informed **(A)** and A*β*-informed **(B)** computational model. PET SUVR values are averaged over all subjects, irrespective of A*β* or tau status. The upper row displays tau seed concentrations mapped onto 3D renderings of the left hemisphere, while the lower row illustrates the corresponding time series. Blue lines represent regions associated with Braak stage 1, and orange lines correspond to Braak stage 2. The anomalous transport scale parameter is *ε* = 5., and the initial concentration of tau seeds *u*_0_ = 0.1 is the same across all regions.

### 3.2. Neuronal activity promotes tau seeding independently of amyloid and tau status

The FDG-based predictions of tau seeding in the medial temporal lobe generalize across subject classifications based on A*β*/tau positivity and is robust to changes in modeling parameters, as shown in

Fig. 3 and Fig. S3 respectively. In particular, we repeat the analysis for A*β*^*−*^, A*β*^+^*τ* MTL^*−*^*τ* NEO^*−*^, A*β*^+^*τ* MTL^+^*τ* NEO^*−*^, and A*β*^+^*τ* MTL^+^*τ* NEO^+^ subjects (see Materials & Methods for more on subject classifications). For each subject classification, we use their averaged FDG PET signals to weight the transport of tau in the model simulations. *The medial temporal lobe is consistently the most vulnerable to seeding across all subject groupings (see Fig. 3) and for varying values of ε (scaling neuronal activity’s impact on axonal transport, see Fig. S3)*.

**Figure 3.**
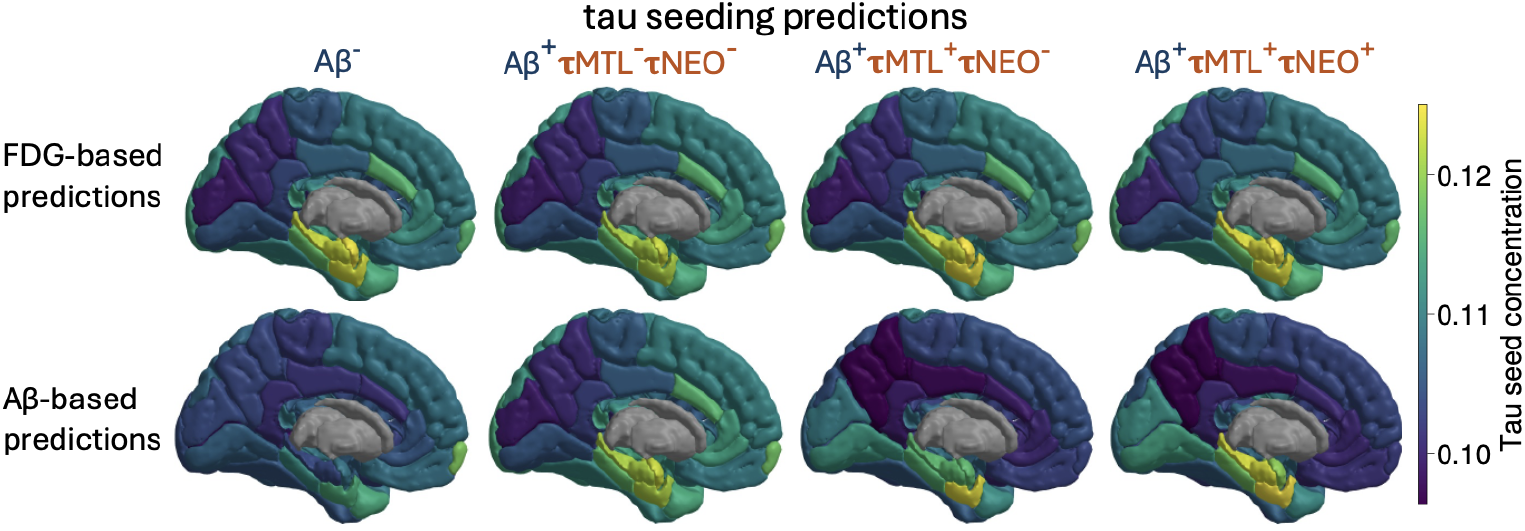
Predicted tau seeding concentrations across subject groups. Predictions of tau seeding given by the computational model informed by FDG- and PET, respectively. The model predictions were informed by PET SUVRs averaged across different subject groups. The tau seed spread throughout the connectome was simulated, and the asymptotic tau seed concentrations are shown on the 3D plots. Columns correspond to subject groups: A*β*^*−*^ (A*β*-negative), A*β*^+^ (A*β*-positive), *τ* MTL (tau deposition in the medial temporal lobe), and *τ* NEO (tau deposition in the neocortex). Rows correspond to FDG- or A*β*-based modeling predictions. The anomalous transport scale parameter is *ε* = 5., and the initial concentration of tau seeds *u*_0_ = 0.1 is the same across all regions.

To further investigate the differences in tau seeding predictions across subject classifications, we computed the distributions in tau seeding predictions in each classification. For each subject in each group, we incorporated their individual FDG PET images to make individualized seeding predictions. We then performed a two-sided Welch’s *t*-test to evaluate the statistical significance of group differences for tau seeding concentrations in Braak stage 1 and 2 regions. Specifically, we tested whether the mean seeding concentrations in the amyloid/tau positive groups differed significantly from the amyloid negative group. As shown in Fig. 4, there is little variation in seeding concentrations between the subject groups in both Braak stage 1 and 2 regions. The A*β*^+^*τ* MTL^*−*^*τ* NEO^*−*^ group is the only group significantly different from the A*β*^*−*^ group and is so only for Braak stage 1 regions.

**Figure 4.**
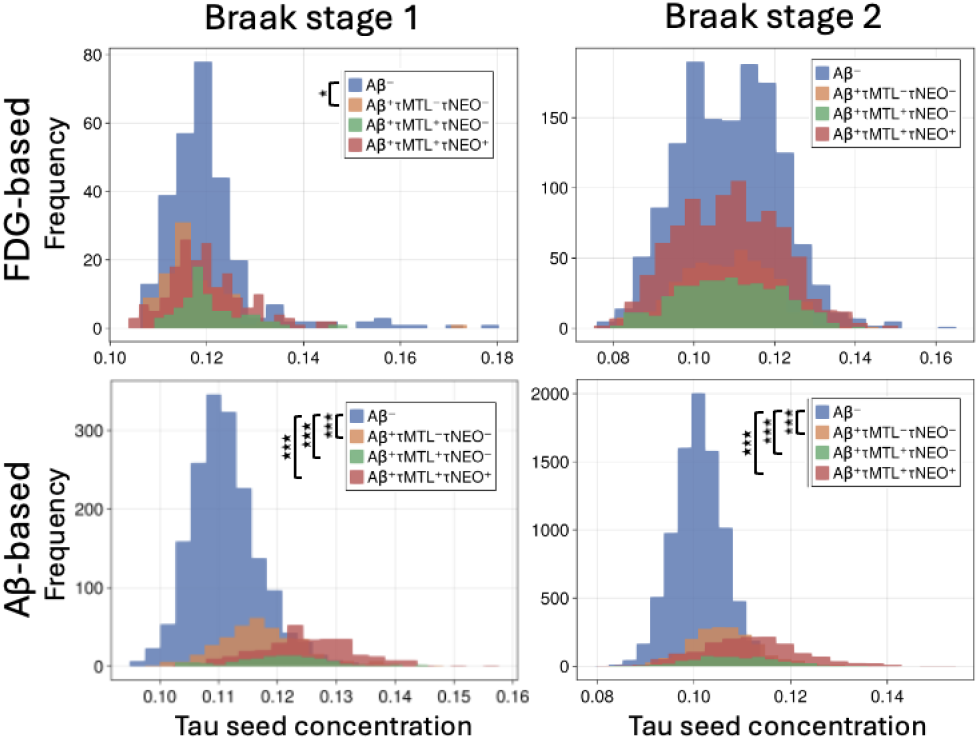
Distributions of predicted tau seed concentrations for subject-level simulations for different subject classifications. The first row shows the distributions for FDG-based simulations, with Braak stage 1 in the first column and Braak stage 2 in the second column. Similarly, the second row presents the distributions for A*β*-based simulations. Each subplot contains four distributions corresponding to different subject classifications. The stars in the legend indicate the results of a two-sided Welch’s *t*-test for differences in average predicted concentrations compared to the A*β*^*−*^ group.

### 3.3. A*β* deposition drives tau seeding in the entorhinal cortex

The medial temporal lobe, and, in particular, the entorhinal cortex, is the first to exhibit tauopathy in Alzheimer’s disease, where, unlike PART, subjects also exhibit A*β* pathology. Furthermore, research suggests that A*β* accelerates the spread of tau pathology,^21,22^ possibly through A*β*-induced hyperexcitability.^23–25^ If this is indeed the case, the presence of A*β* may increase the transport of tau seeds. For this reason, we repeat the simulations and analyses of the previous section, but where A*β* PET signals replace the FDG PET. As shown in Fig. 2B, incorporating the averaged A*β* signal across all A*β*^+^ subjects into the graph Laplacian weighting increases the seeding concentration in the entorhinal cortex. In contrast to simulations where FDG was used to weight tau transport—highlighting the EC, hippocampus, and amygdala as vulnerable regions—A*β* weighting specifically emphasizes the EC as the primary seed region. Moreover, the disparity in seeding concentrations between the EC and the rest of the brain is greater for A*β*-based predictions than for FDG-based predictions, as seen in Fig. 2B.

### 3.4. A*β*-induced tau seeding differs across subject groups

The A*β*-based predictions show greater variation between subject groups (using the average A*β* PET signals from the different subject classifications to inform the model) than the FDG-based predictions, as shown in Fig. 3. Perhaps unsurprisingly, the A*β*^*−*^ group does not exhibit any tau seeding. In contrast, the A*β*-positive groups consistently show greater accumulation in the medial temporal lobe (see Fig. 3), also for varying levels of *ε* controlling the impact of A*β*-deposition on tau transport (see Fig. S4). However, the A*β*^+^*τ* MTL^+^*τ* NEO^*−*^ group (exhibiting tau accumulation in the medial temporal lobe but not in the neocortex) shows a particularly strong bias towards tauopathy in the entorhinal cortex alone. This group has tau and amyloid levels corresponding to early-stage Alzheimer’s disease, thus suggesting that *Aβ-deposition patterns found in early Alzheimer’s are particularly conducive towards tau seeding in the entorhinal cortex*.

Evaluating a two-sided Welch’s *t*-test on the distributions of individualized model-derived tau seed concentrations in early Braak stages across subject groups reveals that A*β*^+^ seeding predictions are significantly different from the A*β*^*−*^ group. As shown in Fig. 4, all A*β*^+^ groups have highly significant differences in their mean seeding predictions for Braak stages 1 and 2 compared to the A*β*^*−*^ group. In particular, the A*β*^+^ groups have higher mean tau seeding levels in early Braak stages, suggesting that amyloid deposition plays a role in the susceptibility of the entorhinal cortex to tau seeding.

### 3.5. Neuronal activity and A*β* are significant predictors of tau seeding in the medial temporal lobe

To test whether the seeding predictions of the computational model are statistically significant, we construct a null model in which the PET signals are shuffled across regions. In this null model, we determine (for each set of parameters and each subject group) a *null concentration threshold*. If a region has a predicted asymptotic concentration above this null threshold, we define it as a *seeding region*. This threshold is the largest gap in all predicted concentrations of the null model over 10,000 simulations with shuffled PET signals (see Materials & Methods for details). The distributions of null model concentrations and the thresholds for the FDG-informed modeling predictions are shown in Fig. S1. Using these thresholds, we perform two statistical tests for each simulation prediction: (i) the probability of observing as many or more Braak stage 1 regions by random chance and (ii) the probability of observing as many or more Braak stage 1 and stage 2 regions. The FDG-informed seeding predictions for each subject group and the significance of each test are shown in Fig. 5A, where we see that each group’s seeding predictions are statistically significant for both tests. For all groups, the left and right entorhinal cortices, hippocampi, and amygdalae pass the null threshold (and are thus seeding regions per our definition). Hence, the predictions that metabolic activity drives the seeding of tau in the medial temporal lobe are significant across all subject groups. As for the A*β*-based modeling predictions, we perform statistical tests in the same fashion as for the FDG-based predictions; see Fig. 5B and S2. For the A*β*-based A*β*-negative group, the stage 1 (entorhinal cortex) seeding predictions are significant, while the stage 2 is not. As for the A*β*-positive groups, all of them have a significant number of stage 1 and stage 2 regions. However, unlike the FDG results, only the entorhinal cortex passes the null threshold in the A*β*-based simulations. Hence, the A*β*-based predictions are more heavily biased towards the entorhinal cortex as opposed to the MTL as a whole. In conclusion, under the assumption that A*β* increases tau transport, brain-wide A*β* patterns significantly predict the entorhinal cortex as the initial tau seeding region.

**Figure 5.**
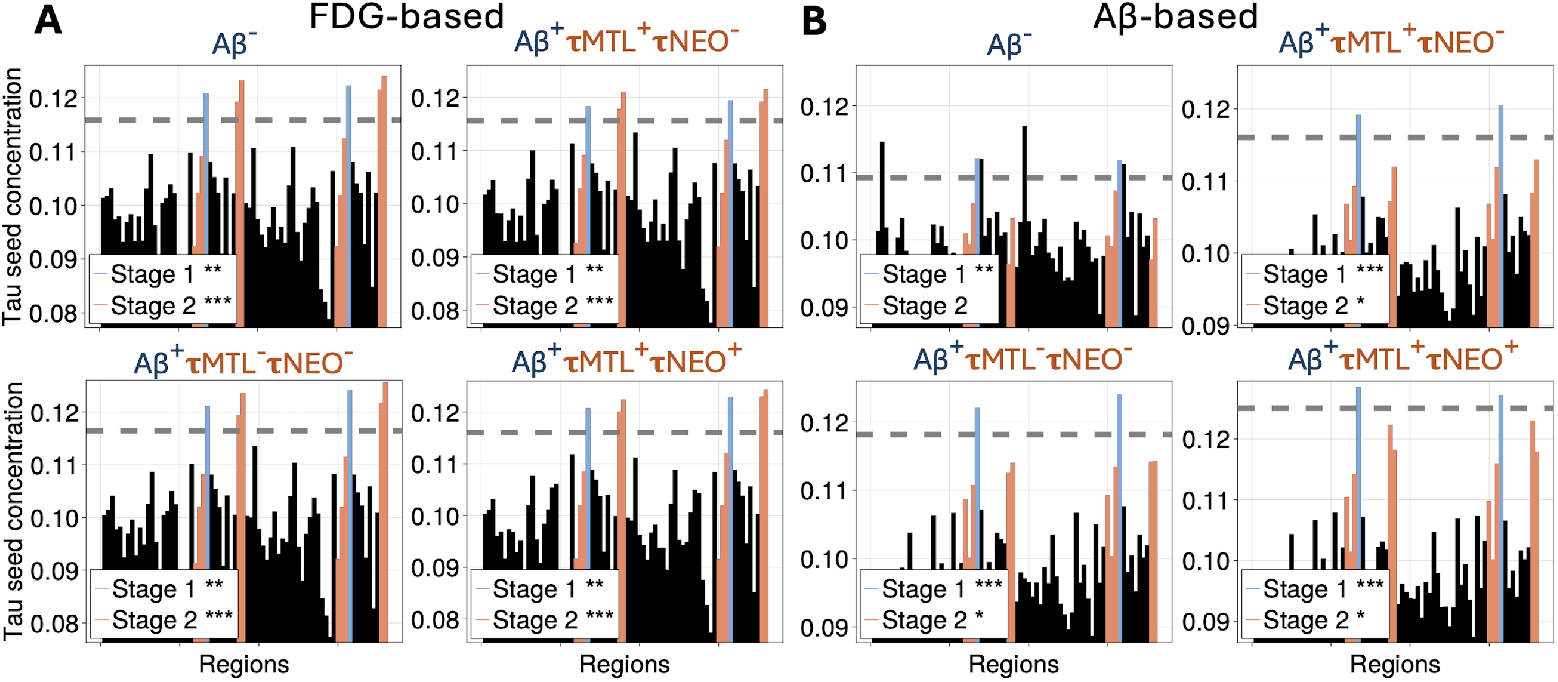
Statistical testing of predicted tau seeding levels across different subject groups. Whole-brain predicted tau seeding levels for the FDG-informed **(A)** and A*β*-informed **(B)** model, with significance testing for early Braak stage seeding.orresponds to A*β*^*−*^ (A*β*-negative), A*β*^+^ (A*β*-positive), *τ* MTL (tau deposition in the medial temporal lobe), and *τ* NEO (tau deposition in the neocortex) and show the asymptotic predicted tau seeds level for each group. The grey stippled line represents the “seeding threshold” determined by the null model (see Materials & Methods). Stars indicate the results of the null hypothesis test, which tests for the probability of observing the same or more stage 1 regions above the seeding threshold and the combined number of stage 1 and 2 above the threshold, when randomly shuffling FDG levels between regions. The anomaly transport scale parameter was set to *ε* = 5

### 3.6. Subject-level neuronal activity and amyloid deposition inform early tau accumulation

In the previous section, we assessed the hypothesis that brain-wide FDG and A*β* PET patterns contribute to early tau seeding in the EC and other MTL regions using subject-averaged PET SUVRs. In this section, however, we assess the hypothesis that FDG and A*β* PET patterns on a subject level contribute to tau seeding in early regions, specifically Braak stage 1 (left and right entorhinal cortices) and 2. To test this, we generate tau seeding predictions for each subject using the same model parameters across all subjects, ensuring that any differences in predictions are solely attributable to the individual FDG and A*β* PET patterns. These predictions are then compared to observed tau SUVR values in the corresponding Braak stages, as shown in Fig. 6. We observe a weak but significant correlation between FDG-based predictions and empirical tau in both Braak stages 1 and 2 (*p <* 0.01, see Materials & Methods for details on regression and statistical analyses). Moderate correlations are observed for A*β*, with highly significant relationships in both stages (*p <* 0.001). These findings lend support to the hypothesis that FDG and A*β* patterns play a role in early tau seeding, as the predicted seeding aligns with empirical tau deposition across subjects.

**Figure 6.**
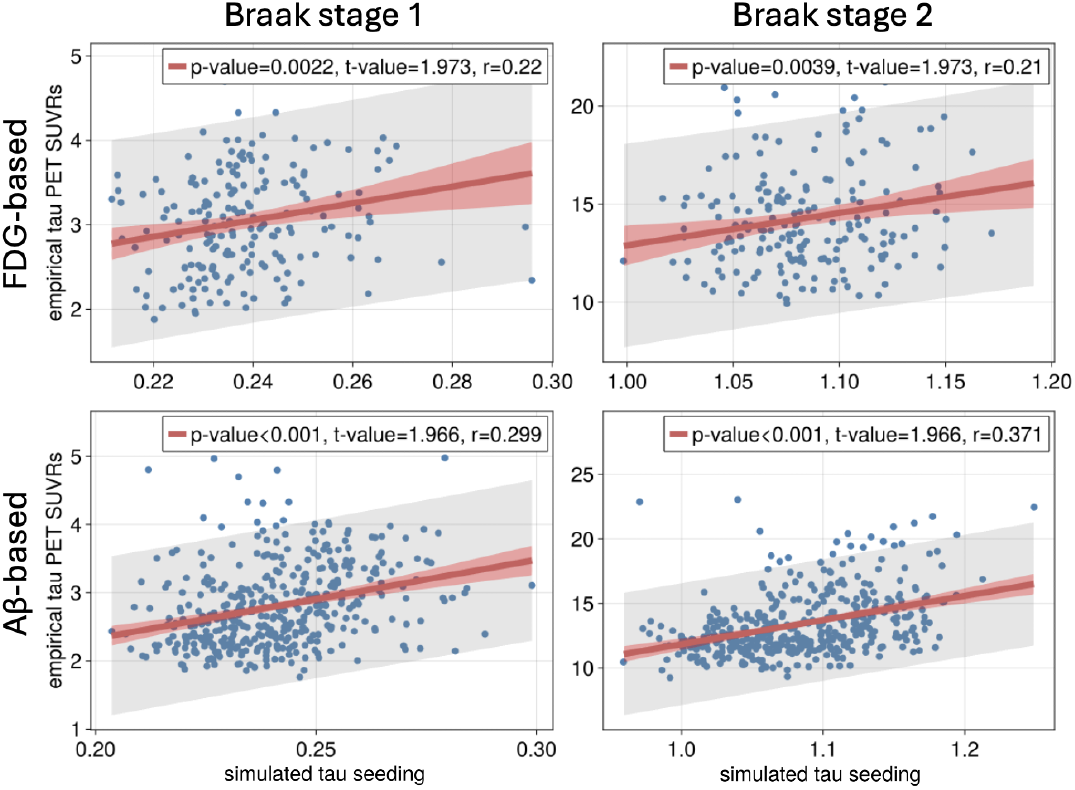
Correlation between model-derived tau seeding and empirical tau PET SUVR in early Braak stages. Each panel shows the comparison between simulated tau seeds and empirical tau SUVRs summed over Braak stage regions, with lines of best fit per least squares linear regression (red line) and 95th-percentile confidence (red shaded region) and prediction intervals (grey shaded region). Legends show the *p*- and *t*-values for the null hypothesis of zero slope and *r*-values (Pearson correlation). The first row displays FDG-based predictions, while the second row presents A*β*-based predictions. The first column compares Braak stage 1 (left and right entorhinal cortex), while the second column compares Braak stage 2. All model simulations used an anomalous transport scaling parameter of *ε* = 5 and an initial seeding value of *u*_0_ = 0.1 for all regions.

## 4. Discussion

The medial temporal lobe, particularly the entorhinal cortex, is consistently the first region to exhibit tau pathology in both Alzheimer’s disease and normal aging. However, the underlying reasons for its heightened susceptibility to tau seeding remain unclear. In this study, we demonstrate that activity-dependent protein transport, where interneuronal tau transport is accelerated by neuronal firing, predicts the seeding of tau in the medial temporal lobe. Specifically, the results support the hypothesis that neuronal activity causes tau seed protein to accumulate in the medial temporal lobe, which would explain the vulnerability of this region to tauopathy in normal aging (PART). Furthermore, assuming that A*β* accelerates tau spread, we show that A*β*-positive individuals additionally exhibit a particular vulnerability to tau seeding in the entorhinal cortex, consistent with early Alzheimer’s pathology. This result suggests that brain-wide patterns of neuronal activity and amyloid deposition may be the underlying force driving tau seeding in the entorhinal cortex.

### 4.1. Contextualizing MTL seeding within whole brain network changes

Recently,^41^ showed with task-based fMRI that A*β* deposition in the default mode network induces a switch from inhibition to activation of the medial temporal lobe and that this directed activation of the MTL correlates with tau accumulation. Broadly speaking, this result corroborates the hypothesis that A*β*-induced hyperactivity leads to tau accumulation in the MTL, which is also supported by our findings. Moreover, our results provide a mechanism for the A*β*-induced EC tau accumulation, where the A*β*-induced hyperexcitability in the default-mode network increases the transport of tau seeds into the MTL. Another recent study by^42^ demonstrated that A*β* deposition is correlated with increased inter-network resting-state fMRI functional connectivity in the limbic, default mode, and control network.^42^ Moreover, increases in functional connectivity between the default-mode and limbic networks correlated with increases in tau PET SUVRs in the limbic network in amyloid-positive, healthy control individuals but not in amyloid-negative, healthy control individuals. Furthermore,^43^ found higher tau accumulation in regions exhibiting A*β*-induced increases in resting-state fMRI functional connectivity to tau epicenters. These findings suggest that A*β* asserts its impact on tau spreading by modulating neuronal communication between brain regions. However, it is unclear how activity-dependent tau transport relates to fMRI functional connectivity metrics, as these measure a notion of synchronicity in neuronal activity between regions and not general neuronal activity levels. These studies establish that functional connectivity changes induced by A*β* may initiate the spread from the MTL to the limbic network and the neocortex. In our study, in contrast, we provide support for the hypothesis that neuronal activity steers tau seeds into the MTL and that A*β* may exacerbate the particular susceptibility of the EC. Although A*β* deposition is necessary for Alzheimer-like propagation of tau in the neocortex, it is not necessary for tau accumulation in the MTL, as is observed in PART. Considering the findings made by^41–43^ along with our own findings, we propose the hypothesis that: (i) brain-wide neuronal activity guides tau seeds to the MTL, regardless of A*β* status, (ii) A*β*-induced hyperexcitability leads to brain-wide changes in neuronal activity and communication, causing the MTL tau to progress to other regions, initiating widespread Alzheimer’s-like tau pathology. Or perhaps, even more simply, amyloid-induced hyperexcitability induces an even higher accumulation in the EC than observed in PART, crossing a critical threshold for which tau starts to spread to the limbic network and eventually the neocortex.

### 4.2. Methodological considerations

While FDG PET provides a robust measure of neuronal activity, it does not capture the full complexity of the neurophysiological processes that may influence tau propagation, such as synaptic activity or intracellular signaling pathways, and may be complemented with fMRI, MEG, EEG and transcriptomic data in future studies. We have also assumed that neuronal activity remains unchanged through time, whilst in reality, neuronal activity itself is a dynamical process evolving over time. Previous models have investigated the co-evolving dynamics of neuronal activity and protein spreading and may be used forward to test further hypotheses on the activity-spreading link.^29,44^ Our results are limited by the lack of directedness in structural connectivity, for which no consensus inference method exists for human imaging. Without directionality in the connectome edges, the structural connectivity does not impact the steady-state tau accumulation in each region (though it does impact the temporal spreading patterns). Nonetheless, our results show that the effect of including the anomalous transport process (neuronal activity and amyloid-*β*) biases the accumulation of tau towards the MTL and EC, regardless of the topology of the structural connectome. This is because the asymptotic tau concentration in each region *i* scales with 1*/*(1 + *εA*_*i*_) independently of the underlying structural connectivity (see Section 2.2). Future studies may explore methods to infer the directionality of structural connectivity to probe the interaction between structure and neuronal activity on regional tau pathology levels. Moreover, our model assumes a linear relationship between neuronal activity, A*β*, and tau transport. In reality, the interaction between these elements may be nonlinear or context-dependent. Furthermore, it should be possible to expand our computational model to include non-linear effects mimicking regional prion-like replication.^45^ Such a model can then simulate the initiation of the disease in the MTL in addition to the ensuing spatiotemporal Braak staging pattern without the need to artificially instruct the computational model to begin in the MTL. As such, computational modeling of prion-like spreading may not only predict individual spreading patterns but also identify the brain regions where propagation is most likely to initiate. Indeed, it has been established that there is considerable variation in the localization of tau pathology between individuals with Alzheimer’s disease,^46^ raising the question whether this variability may be, in part, accounted for by differences in brain-wide neuronal activity and A*β* deposition.

### 4.3. Conclusion

In this study, by assuming that neuronal firing and amyloid-*β* accelerate cell-to-cell transport of tau, we demonstrated that brain-wide patterns of neuronal activity and amyloid-*β* explain the susceptibility of the MTL to tau seeding, particularly in the EC, in Alzheimer’s disease and PART. Moreover, our findings imply that increased neuronal activity in the medial temporal lobe will lower its susceptibility toward tau seeding. Anecdotally, taxi and ambulance drivers are the least likely to die from Alzheimer’s disease among all occupations,^47^ which is hypothesized to be related to functional changes in the hippocampus, as observed in London taxi drivers.^48^

Our model suggests a mechanism for this hypothesis, in which the increased hippocampus activation may clear accumulating tau away and re-distribute the tau seeds to non-pathological levels in surrounding regions. Our findings propose a “*use it or lose it* “principle, in which regions with low neuronal activity levels experience higher chances of tau pathology. As such, one may hypothesize that neuronal stimulation of structures in the medial temporal lobe can reduce tau accumulation. Indeed, deep brain stimulation of the entorhinal cortex in mouse models of AD has shown promise in decreasing tau levels and ameliorating cognitive symptoms.^49,50^ Further research is needed to clarify the role of neuronal activity on tau transport, the role of A*β* in tau seeding, and potential therapeutic strategies based on neuronal stimulation.

