## Supplemental information for "Neuronal activity and amyloid-β cause tau seeding in the entorhinal cortex in Alzheimer’s disease"

### Null model for seeding susceptibility

As described in Materials & Methods, we construct a null model for seeding predictions, where the computational model is integrated with PET SUVRs randomly shuffled across regions. To evaluate the probability of observing as many or more seeded regions belonging to Braak stage 1 and 2 under the null hypothesis, we shuffle the PET SUVRs and collect all the asymptotic tau concentrations across all regions into one dataset, providing us with a distribution of null concentrations. These distributions, computed for each subject group, are shown in Figures S1 and S2 for the FDG- and A$\beta$-based models respectively. These distributions of concentrations reveal the distributions of predicted concentrations upon randomly shuffling PET SUVRs. We then define the *seeding threshold* as the midpoint in the largest gap between any two values in these distributions, shown as a grey stippled line in Figures S1 and S2. To test the significance of our seeding predictions, we consider a region with predicted concentrations beyond the seeding threshold to be seeded, and compute then the likelihood of observing as many Braak stage 1 (and 2) regions as seeded in the null model, which serves as the $p$-value.

### Varying the impact of neuronal activity and amyloid on modeling predictions

By varying the control parameter $\varepsilon$, we adjust the impact that neuronal activity and amyloid has on the predicted seeding susceptibility across the brain. As seen in Figures S3 and S4, for low levels of $\varepsilon$

*Data used in preparation of this article were obtained from the Alzheimer's Disease Neuroimaging Initiative (ADNI) database (adni.loni.usc.edu). As such, the investigators within the ADNI contributed to the design and implementation of ADNI and/or provided data but did not participate in the analysis or writing of this http://adni.loni.usc.edu/wp-content/uploads/how_to_apply/ADNI_Acknowledgement_List.pdf

§These authors contributed equally to this work
✉

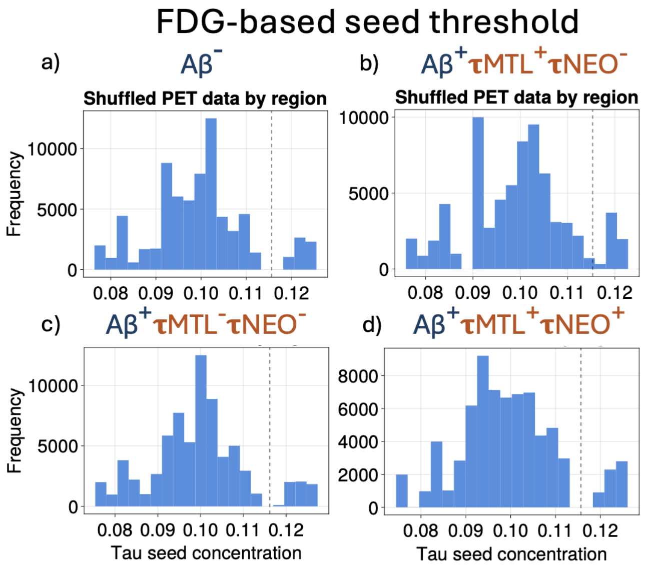


Figure S1: Histograms of seeding concentrations pooled across all brain regions, with seeding concentrations calculated for both the original and shuffled FDG PET signals. The shuffled PET signals are generated by randomizing the spatial distribution of PET values across regions. The seeding threshold, determined by the maximum difference method, is shown as a grey stippled line to highlight the threshold for significant seeding concentrations.

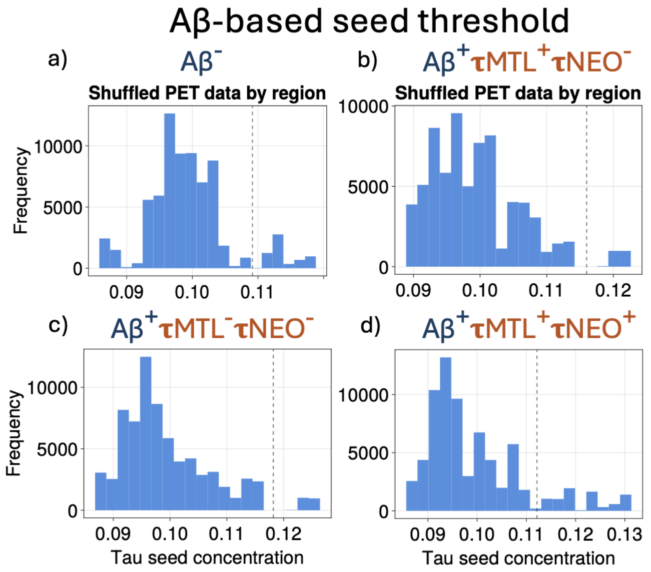


Figure S2: Histograms of seeding concentrations pooled across all brain regions, with seeding concentrations calculated for both the original and shuffled A$\beta$ PET signals. The shuffled PET signals are generated by randomizing the spatial distribution of PET values across regions. The seeding threshold, determined by the maximum difference method, is shown as a grey stippled line to highlight the threshold for significant seeding concentrations.

there is no bias in seeding towards any particular region, and as $\varepsilon$ is increased these biases become more apparent.

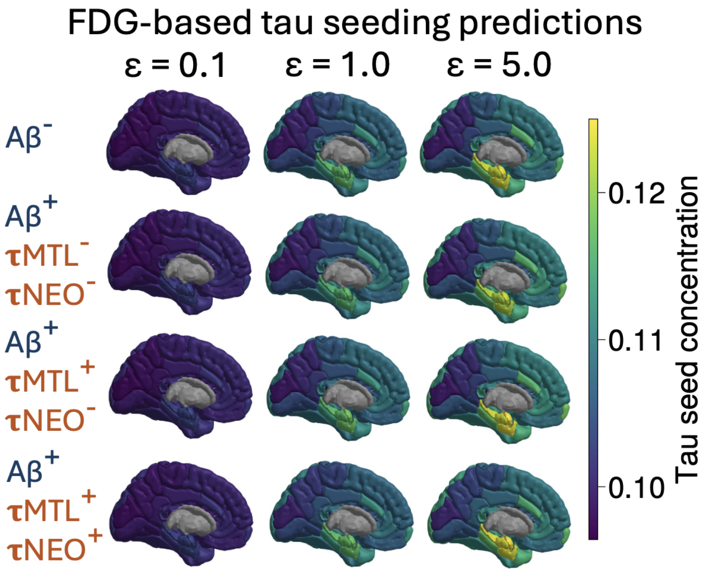


Figure S3: **Predicted tau seeding concentrations across subject groups and FDG impact levels.** Rows correspond to subject groups: $A\beta^-$($A\beta$-negative), $A\beta^+$($A\beta$-positive), $\tau$MTL (tau deposition in the medial temporal lobe), and $\tau$NEO (tau deposition in the neocortex). Columns represent increasing levels of the predicted impact of metabolic activity (FDG) on tau seeding, progressing from left to right.

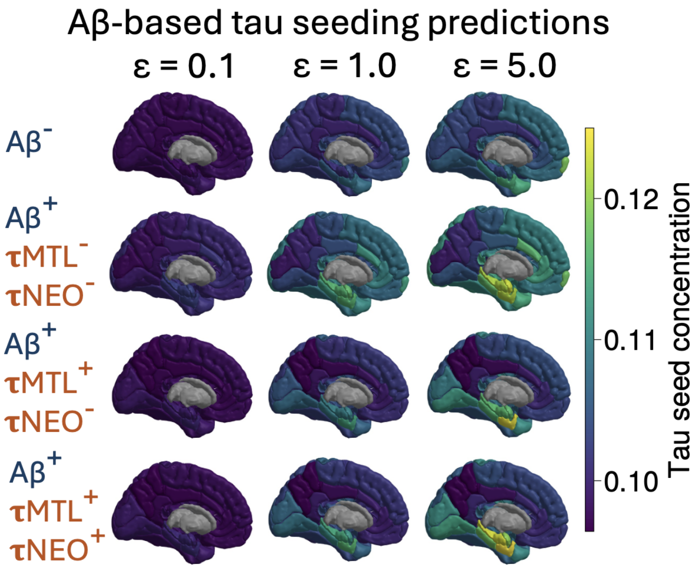


Figure S4: **Predicted tau seeding concentrations across subject groups and A$\beta$ impact levels.** Rows correspond to subject groups: $A\beta^-$($A\beta$-negative), $A\beta^+$($A\beta$-positive), $\tau$MTL (tau deposition in the medial temporal lobe), and $\tau$NEO (tau deposition in the neocortex). Columns represent increasing levels of the predicted impact of A$\beta$-deposition on tau seeding, progressing from left to right.
